# Fluorophore-labelled RNA aptamers to common protein tags as super-resolution imaging reagents

**DOI:** 10.1101/2020.02.27.968578

**Authors:** Juan Wang, Avtar Singh, Abdullah Ozer, Warren R. Zipfel

## Abstract

Developing labelling methods that densely and specifically label targeted cellular structures is critically important for centroid localization-based super-resolution microscopy. Being easy and inexpensive to produce in the laboratory and of relatively small size, RNA aptamers have potential as a substitute for conventional antibody labelling. By using aptamers selected against common protein tags - GFP (green fluorescent protein) in this case - we demonstrate labelling methods using dSTORM-compatible fluorophores for STORM and hybridizable imager strands for DNA-PAINT super-resolution optical imaging of any cellular proteins fused to the aptamer binding target. We show that we can label both extracellular and intracellular proteins for super-resolution imaging, and that the method in particular, offers some interesting advantages for live cell super-resolution imaging of plasma membrane proteins.

**KEY POINTS:**

1. A simple to use RNA aptamer method for super-resolution STORM and PAINT imaging in cells expressing common protein tags.
2. The method has a number of unique advantages for live cell imaging at the nanometer scale.
3. Provides a means to quantify the number of proteins being trafficked on the plasma membrane.

## INTRODUCTION

Optical microscopy is a mainstay of nearly all biological and biomedical studies. However, for many applications a major limitation is the optical resolution since many cellular structures are below the diffraction limit of ~250 nm making them un-resolvable using conventional microscopy. The past decade has seen the development of a number super-resolution (SR) imaging techniques that circumvent the diffraction limit. The three main super-resolution techniques consist of centroid localization-based modes such as Photo-Activated Localization Microscopy (PALM) (1) and Stochastic Optical Reconstruction Microscopy (STORM) (2), Stimulated Emission Depletion (STED) microscopy (3) and super-resolution Structured Illumination Microscopy (SR-SIM) (4). Centroid-based SR imaging methods work with densely labelled samples by either activating only a small subset of the fluorophores (PALM) or turning most of them “off” by conversion to a dark state (STORM). STED microscopy uses an engineered depletion beam point-spread function that deactivates fluorophores outside of the center of the excitation laser focus. The resolution achievable with STED image is dictated by the shape and scale of the STED beam focus. SIM achieves super-resolution by collecting spatial frequencies beyond the conventional range of the objective lens by illuminating the object with patterned light. The resolution improvement in SR-SIM is limited to a factor of two, while PALM/STORM or STED methods can achieve a ~10-fold increase in resolution.

Here we focus primarily on developing a system of unique probes for applications of STORM; however, our method is a widely applicable labelling technique for all modes of fluorescence microscopy. There are currently several labelling techniques used in super-resolution imaging. Two most common methods involve tagging the protein of interest (POI) with photoactivatable fluorescent proteins such as PA-GFP or mEOS (5), or enzymatic tags like SNAP- or HALO-tags that covalently react with cell-permeable organic dyes (6,7). These methods require the creation of cells stably expressing fusion proteins with these tags or transient transfection of the protein of interest (POI) Genetically encodable photoactivatable fluorescent proteins offer a clear advantage for studies of cell dynamics, enabling live-cell imaging. However, many of the available fluorescent proteins have relatively poor photo-stability and low molecular brightness compared to organic dyes. Although organic dyes for conventional live cell imaging in the form of indicators and organelle specific dyes are commonly used, direct targeting of specific proteins by organic dyes, as needed for most super-resolution imaging experiments, is more difficult. Cells expressing modified target proteins with enzymatic tags such as SNAP and CLIP, which bind and activate a modified cell-permeable organic fluorophore can be used. This helps overcome the photostability and molecular brightness issues common with many photoactivatable fluorescent proteins, but the intracellular labelling efficiency and uniformity can be low with these tags. For permeabilized and fixed cells, antibodies labelled with fluorescent dyes have long been used for all modes of fluorescence microscopy and are the main labelling method for STORM and STED imaging.

Another class of molecules that have recently shown promise are aptamers. They are either small nucleic acid oligos or short peptides (8, 9) selected based on tight binding to specific molecular targets. Nucleic acid-based aptamers are generated through an in vitro process called Systematic Evolution of Ligands by Exponential Enrichment (SELEX) (10,11) where a pool of random nucleic acid sequences undergoes a number of selection rounds, so that sequences with high affinity and specificity for the target molecule are selected. RNA aptamers can have high binding affinities, are small (2-3 nm vs 10-15 nm for antibodies), and are easy and inexpensive to produce in the laboratory, making them a useful substitute for conventional antibody labelling (12). A caveat of using aptamers for labelling proteins of interest is that it is tedious to select and verify the binding specificity of an aptamer to its target protein. Therefore, a more generic super-resolution labelling method that uses aptamers against common tag proteins would be beneficial since it avoids the need to carry out SELEX for each new POI and laborious characterization of binding specificity. This strategy would enable studies with cell lines expressing tagged POIs either stably or transiently. Since a multitude of cell lines with GFP tagged proteins of interest and a well-characterized, high affinity RNA aptamer is already available, we have initially focused our efforts on using the GFP-binding aptamer (AP3) (13,14) for our proof-of-concept aptamer-POI labeling strategy for high-resolution microscopy. AP3 tightly binds GFP and EGFP (Kd ~4 nM) and reduces its fluorescence under 488 nm excitation at physiological pH by increasing the pKa of EGFP from 5.8 to 6.7. AP3 also binds to GFP derivatives YFP and CFP with a slightly lower affinity (~10 nM Kd). The change in GFP pKa indicates that AP3 interacts with the protein in a way that modifies the barrel structure allowing easier access of protons to the chromophore protonation sites.

Here we introduce a method of targeting GFP-labelled proteins for direct stochastic optical reconstruction microscopy (dSTORM)-based super-resolution imaging using Alexa-647 labelled AP3 aptamer. Because aptamers are not readily cell permeable, previous super-resolution imaging done with labelled aptamers were restricted to membrane proteins (12). We have optimized our labelling protocol to extend this method for super-resolution imaging of intracellular proteins and demonstrate this on cells stably expressing EGFP-tagged Lamin A, a nuclear protein localized to surface of the nuclear envelope and associated with heterochromatin. We also show that we can apply our labelling method for live-cell super-resolution imaging and for PAINT (Points Accumulation for Imaging in Nanoscale Topography) based super-resolution imaging, a method that relies on the transient binding of fluorophores to the target from a diffusing pool of labelled DNA imager strands (15, 16). The use of aptamers for PAINT SR application greatly simplifies the method. Overall, we show that protein tag-targeting RNA aptamers are a powerful substitute for antibody labelling for a number of modern fluorescence imaging modalities.

## MATERIAL AND METHODS

### Sequence of AP3, AP3-docking and Alexa 647 labelled imager strand

GFP aptamer (AP3) sequence: AGCTTCTGGACTGCGATGGGAGCACGAAACGTCGTGGCGCAATTGGGTGGGGAAAGTCCTTAAAAGAGGGCCACCACAGAAGCT

T7pro-AP3 Forward primer (T7 RNA Polymerase promoter used for in vitro transcription of the RNA aptamer is underlined): <u>GAA TTA ATA CGA C</u>TC ACT ATA GGG AGCT TCT GGA CTG CGA TGG GAG CA

AP3 Reverse primer: GCT TCT GTG GTG GCC CTC TTT TAA GGA CT

AP3 Reverse primer with 9nt docking site (underlined): <u>TAGATGTAT</u>ATAAAGCTTCTGTGGTGGCCCTCTTTTAAGGACT

Imager strand: CTAGATGTAT-Alexa 647

All oligos were purchased from IDT with standard desalting and used without any further purification.

#### *In vitro* transcription of GFP aptamers

GFP aptamer DNA templates were generated by PCR and purified by 8% native PAGE gel extraction. *In vitro* transcription reaction with T7 RNA polymerase was incubated at 37°C overnight, followed by phenol-chloroform extraction and isopropanol precipitation. The precipitated GFP RNA aptamers were re-suspended in DEPC-treated ddH_2_O and un-reacted free nucleotides were removed up with two sequential Micro Bio-spin P-30 columns (RNase-free).

#### AP3-Alexa 647 3’ end labelling chemistry

GFP aptamers from in-vitro transcription prep were oxidized with 0.5 mM NaIO_4_ in 100 mM NaOAc (pH 5.1) at room temperature for at least 90 minutes. They were then precipitated with ice-cold ethanol and re-suspended in 100 mM NaOAc buffer. The oxidized GFP aptamers were incubated with Alexa 647 hydrazide (Thermo-Fisher Scientific Cat #: A-20502) at 1:10 ratio in the fridge (4°C) overnight. The labelling reaction was purified the next day with two Micro Bio-spin P-30 columns (RNase-free) to remove excessive unreacted dyes.

#### Comparison of labeled vs unlabeled GFP aptamer binding in solution

GFP aptamers were incubated with 50 nM GFP at various aptamer concentrations in aptamer binding buffer (PBS, 5 mM MgCl_2_, pH 7.4) for 5 min at room temperature and emission spectra were measured using a model QM4 fluorometer (Photon Technology International/Horiba, Birmingham, NJ). The emission spectra (490 - 600 nm) of AP3 bound GFP at various concentrations were acquired using 480 nm excitation. The same measurements were repeated with Alexa 647 labelled GFP aptamers. Dissociation constants (Kd) were determined for plots of 1-F/Fo vs aptamer concentration and data fit to the Morrison equation (17) to determine the Kd.

#### Preparation of fluorescent protein (FP) immobilized Ni-NTA beads

Ni-NTA beads were washed 3X with 10 volumes of washing buffer (20 mM Tris.Cl, 125 mM NaCl and 25 mM KCl, pH 7.5) and re-suspended at 20% v/v in the washing buffer. His-GFP, His-mNG, or His-mCherry were then added (1 μg of His-tagged FP to 1ul of beads) to the bead solution and incubated at 4° C on a rotator for an hour. After the binding of His-FP to the beads, unbound His-FPs were removed by spinning down and re-suspending in 10 volumes of wash buffer for 3 times. The FP bound Ni-NTA was kept in washing buffer at 4°C at most for a few days before experiments.

#### Fibronectin Treatment of coverslips

Coverslip bottom dishes (MatTek Corporation, Ashland, MA) were treated with 1M NaOH for 30 min, then washed 3X with autoclaved water and incubated in 4% MPTS in EtOH for 30 min. Next, they were washed 3X with 100% EtOH and coated with 4 mM GMBS in EtOH for 30 min. Afterwards they were washed gain twice with 100% EtOH and then twice with PBS. Finally, dishes were coated with 10μg/ml fibronectin solution (Sigma-Aldrich, F1141) at 4°C overnight.

#### Cell culture and fixation for STORM imaging of TfR-EGFP

HEK293 cells stably expressing TfR-GFP were plated on fibronectin coated MatTek dishes and the next day incubated with 1 μM Alexa 647 labelled AP3 aptamer in blocking buffer (2 mM salmon sperm DNA, ThermoFisher #15632011), 2 mM yeast tRNA (ThermoFisher #AM7119) at room temperature for at least 20 min. The cells were washed three times with GFP aptamer binding buffer and fixed with 4% PFA in PBS for 10 min. **Lamin A-EGFP cells**: Human fibroblast cells stably expressing Lamin A-EGFP (kindly provided by Jan Lammerding, Cornell University) were plated on fibronectin coated MatTek dishes. The next day, cells were permeabilized with 0.2% Triton X-100 once in “buffer S” (4% PEG 8000, 1mM EGTA, 0.1M PIPES in DEPC-treated ddH2O) for 4 min. Cells were then incubated with 1 μM Alexa 647 labelled GFP aptamer in aptamer binding buffer at room temperature for 20 min, washed three times with aptamer binding buffer and fixed with 4% PFA for 10 min at room temperature.

#### Aptamer PAINT for Lamin A-EGFP cells

Human Fibroblast cells stably expressing Lamin A-EGFP were plated on fibronectin coated MatTek dishes a day before staining. When the cells were ~90% confluent they were washed twice with PBS and permeabilized with 0.2% Triton X-100 in buffer S for 4 min. Next, the cells were incubated with aptamer staining solution (1.5μM AP3-P1-docking, 2 mM ytRNA, 2 mM salmon sperm DNA) for 20 min and fixed with 4% PFA at room temperature for 20 min before imaging. **TfR-EGFP cells:** HEK293 TfR-EGFP cells were plated on the fibronectin coated MatTek dishes a day before imaging. The cells were ~90% confluent when stained with AP3-P1-docking. The dish was washed twice with PBS and then incubated with aptamer staining solution for at least 1h. The staining solution contained 1.5 μM of AP3-P1-docking, 2mM ytRNA, 2 mM salmon sperm DNA.

##### PAINT Imaging

Imager strands were added to the solution at a final concentration of 7 nM and imaged on a Zeiss Elyra. PS.1 microscope as described below.

#### Anti-GFP antibody-Alexa 647 Staining of HEK293-EGFP cells

HEK293 EGFP cells were plated on non-coated MatTek dishes a day before and fixed with 4% PFA in 1x PBS for 15 min at RT. Cells were washed twice with PBS for a 5 min incubation period each time. After the PBS, wash the sample was blocked by incubation with 5% BSA in PBS for 30 min, and then stained with 10 μg/ml Alexa 647 labelled Anti-GFP antibody for 1 hour. Finally, cells were washed twice with 1% Tween-20 in PBS for 5 min each time and stored in PBS for imaging.

#### Phalloidin-Alexa 647 Staining of HEK293-EGFP cells

HEK293 EGFP cells plated on non-coated MatTek dishes the day before were fixed with 4% PFA in PBS for at least 10 min at RT. The cells were washed twice with PBS at RT for 5 min each time. 0.5% Triton-X100 solution was added to the dish to permeabilize the cells for 10 min at RT. After a PBS wash the sample was blocked by incubation with 5% BSA for 30 min, stained with Phalloidin-Alexa 647 (200:1 dilution) for 1 hour, and then subjected to two 5 minute washes with 1% Tween-20 in PBS and stored in PBS for imaging.

### Microscope Setup and Imaging Analysis

#### Confocal images of AP3-Alexa 647 binding to FP immobilized beads

Images were acquired on a Zeiss LSM880 confocal system with a 40x Plan-Apochromat, oil immersion, 1.4 NA objective. EGFP and mNG immobilized beads bound by AP3-Alexa 647 were excited using 488 nm and 632 nm laser lines, respectively. mCherry immobilized beads bound by AP3-Alexa 647 were imaged using 561 nm and 632 nm laser excitation.

#### STORM and PAINT images of HEK 293 TfR-EGFP cells and Human Fibroblast Lamin A-EGFP cells

STORM and DNA-PAINT images were acquired on a Zeiss Elyra PS.1 super-resolution system with a Zeiss 100x Plan-Apochromat oil immersion, 1.46NA TIRF objective. STORM images of HEK293 TfR-EGFP cells and Human Fibroblast Lamin A-EGFP cells were acquired at a frame rate of 50 Hz for 50,000 frames. PAINT images of fixed Human Fibroblast Lamin A-EGFP cells and live HEK293 TfR-EGFP cells were acquired at a frame rate of 20 Hz and 100 Hz, respectively. PAINT and STORM images were analyzed and reconstructed with Zeiss Elyra PS.1 Zen analysis software.

## RESULTS

### *In vitro* binding test of AP3-Alexa 647 - binding affinity and specificity

For dSTORM super-resolution imaging (15) we covalently labeled AP3 aptamer with Alexa 647 dye at its 3’-end. To test whether the Alexa 647 label altered the binding affinity of AP3, we measured the disassociation constant Kd of both the unlabeled and labeled aptamers in solution and found that the added fluorophore did not change the binding affinity. The Kd’s of the labeled and unlabeled aptamers were both in the 4 nM range (Fig. 1a, Supplementary Fig. S1a), with the Kd of unlabeled AP3 aptamer in good agreement with our previously published result (13, 14). We also tested the binding of Alexa 647 labeled AP3 on EGFP immobilized on Ni-NTA beads (Fig. 1b). EGFP-bound beads incubated with AP3-Alexa 647 showed strong fluorescence in red channel due to binding of AP3-Alexa 647 to the EGFP coated beads, with no detectable signal for the three control conditions. To determine the binding specificity of the AP3 to EGFP we used Ni-NTA beads bound with non-GFP derived fluorescent proteins and found that AP3 does not bind to mNeonGreen and mCherry (Supplementary Fig. S1b) indicating that AP3 is highly specific only to EGFP and its derivatives. We also found that in general, AP3 does not bind PFA fixed GFP or EGFP to any useable extent (Supplementary Fig. S2), presumably due to cross-links formed between GFP and neighboring proteins that block the AP3 binding interaction, or modification of surface lysine residues critical for the binding of AP3 aptamer. Similar to antibodies, aptamers only optimally bind targets in the state that they were selected against. If aptamers binding to fixed proteins are desired, it may be selections could be made against fixed targets to isolate tight binding aptamers.

**Figure 1.**
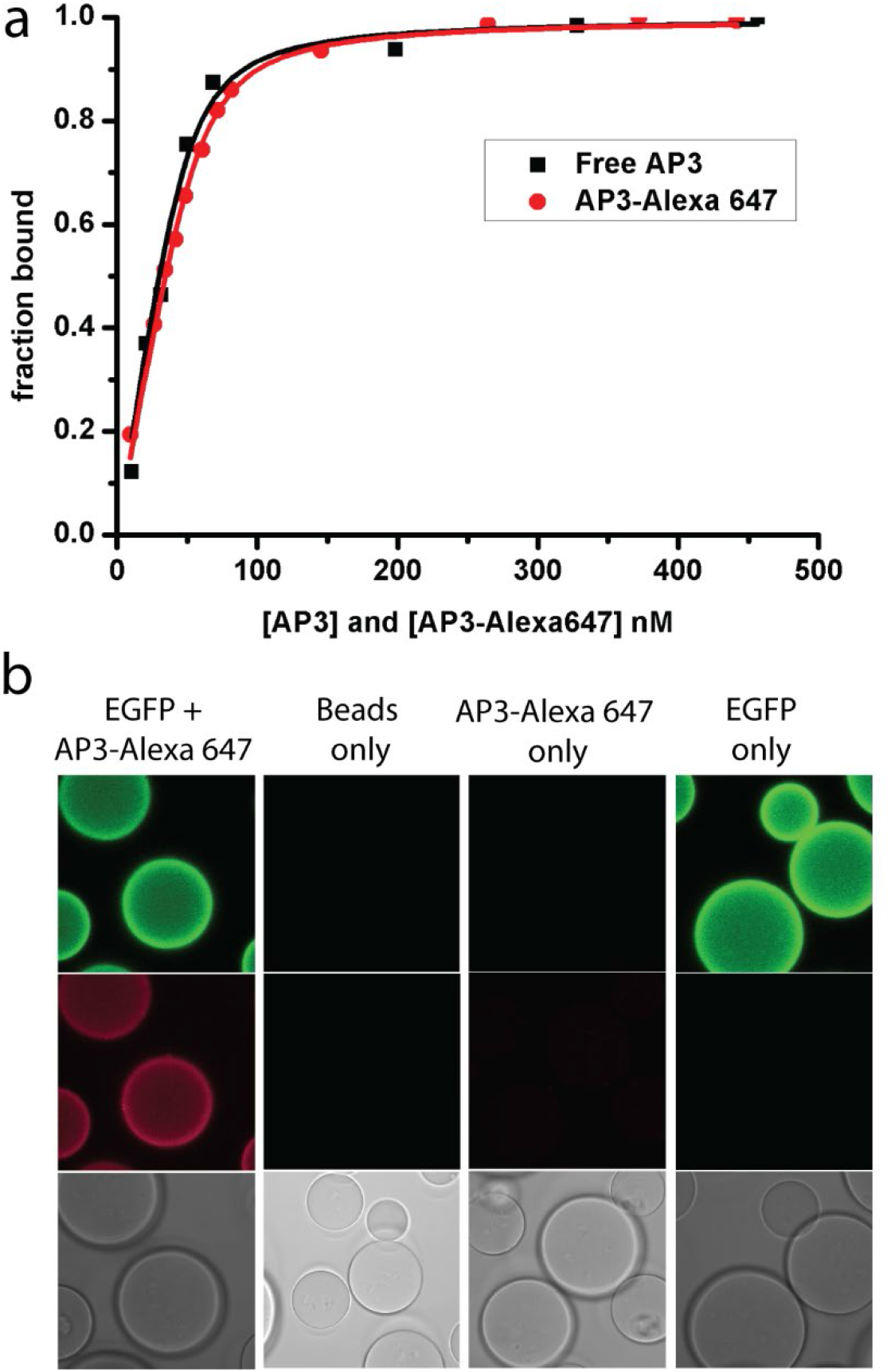
*in vitro* binding characterization of AP3-Alexa 647. (a) The binding affinity of free AP3 and AP3-Alexa 647 to EGFP in solution was measured based on the decrease in EGFP signal caused by aptamer binding. Data was plotted a 1-F to represent the fraction bound and the curves to the quadratic binding equation. The Kd for free AP3 and Alexa 647 labeled AP3 were both determined to be < 5 nM, indicating that the Alexa 647 label does not change the binding affinity of AP3. (b) AP3-Alexa 647 binding to immobilized EGFP on Ni-NTA beads. The right three panels are negative controls.

### Conventional and super-resolution imaging of AP3-Alexa 647 binding to GFP tagged cellular structures

We verified that AP3-Alexa 647 binds extracellular proteins on the surface of cells using HEK293 cells that stably express transferrin receptor-EGFP fusion protein, TfR-EGFP. TfR is transported to the plasma membrane where it binds transferrin, forms clusters on the cell membrane, and then are internalized via the endocytosis process leading to internalization of transferrin. After incubation with our fluorescent aptamer probe, we imaged AP3-Alexa 647 stained cells using conventional microscopy (Fig. 2a). Total internal refection microscopy (TIRFM) images (Fig. 2b-d) show colocalized signals from a subset of the TfR-EGFP (Fig. 2b). TIRF illumination only penetrates a few hundred nanometers into the cell near the coverslip surface and the high concentration of TfR-EGFP near the inner leaflet of the membrane generates a higher background in the green channel since the internalized TfR-EGFPs are not accessible to the AP3-Alexa 647.

**Figure 2.**
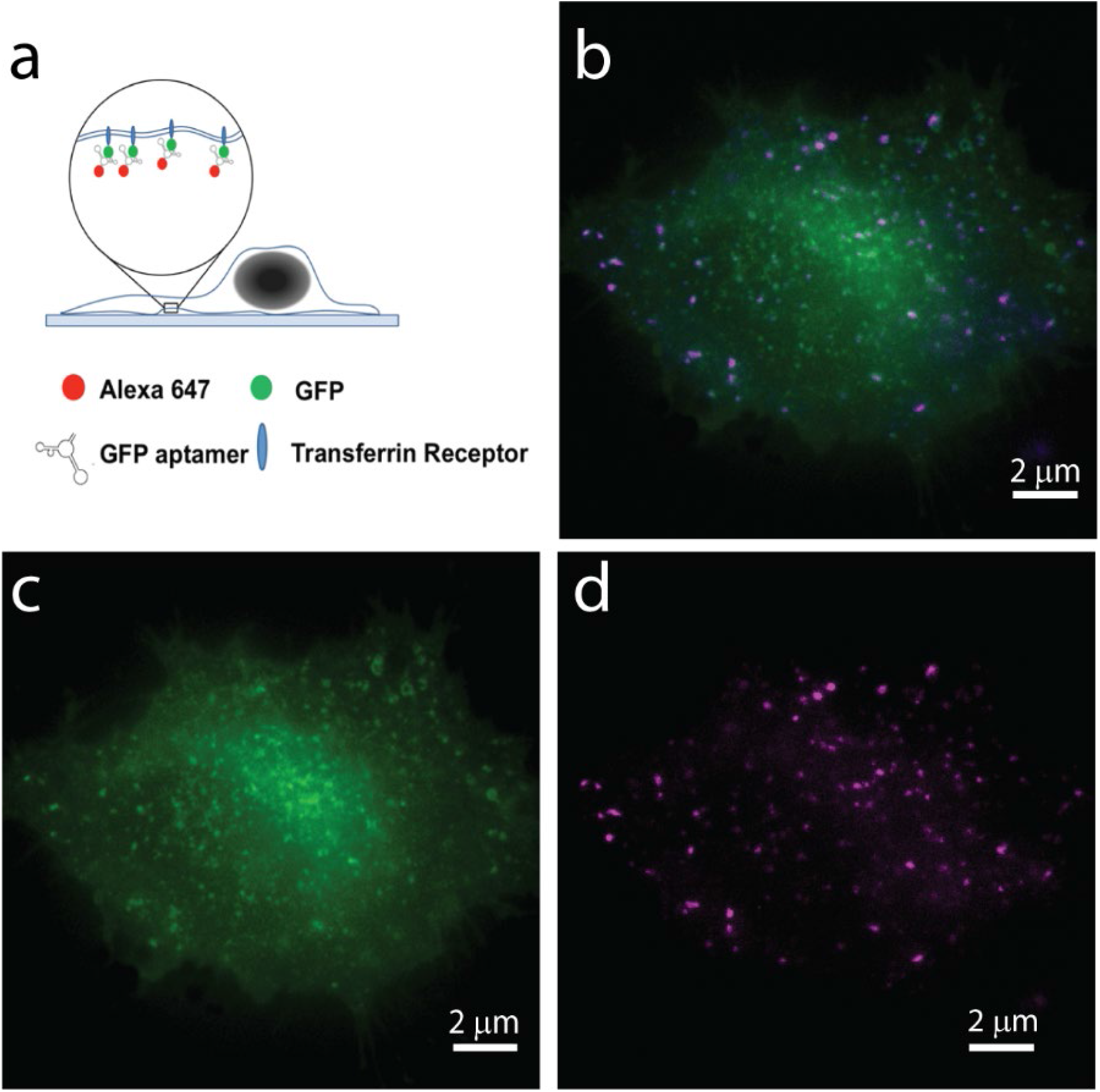
TIRFM images of AP3-Alexa 647 labelled TfR-EGFP expressing cells. (a) AP3-Alexa 647 is cell impermeable and only labels TfR-EGFP on the outer membrane. (b) Merged image (green: EGFP, magenta: Alexa 647). (c) EGFP channel - higher background in the green channel is due to internalized TfR-EGFP near the inner leaflet. (d) Extracellularly exposed TfR-EGFP labelled with AP3-Alexa 647.

TfR is known to form sub-diffraction ring structures as it clusters around an endocytic pore on the plasma membrane, making it an ideal system for testing our aptamer labeling method for super-resolution imaging. HEK293 cells expressing TfR-EGFP were paraformaldehyde fixed and imaged using the standard dSTORM protocol using AP3-Alexa 647 for labeling. dSTORM images were acquired using a Zeiss Elyra PS.1 microscope at a frame rate of 50Hz for 50000 frames and reconstructed using the Zeiss software. Lateral drifts were corrected using 100 nm TetraSpeck beads as the fiducial markers. Figure 3a shows the TIRF (EGFP channel) and a dSTORM reconstruction (Alexa 647 channel) of the same cell. Figure 3b shows the boxed region in 3a demonstrating the improved resolution attained using the AP3-Alexa 647 and dSTORM. Compared to TIRF, the dSTORM image has much higher resolution and shows distinct shapes of the transferrin receptor clusters. Analysis shows that we can distinguish two transferrin receptors (indicated by the yellow arrow) that were ~70nm apart on the dSTORM reconstruction (Fig. 3c). We measured the size distribution of TfR clusters using reconstructed dSTORM images of TfR-EGFP cells (Fig. 3d and Supplementary Fig. S3) and found an average cluster size (~80 nm) that corresponds with other super-resolution microscopy results reported (16).

**Figure 3.**
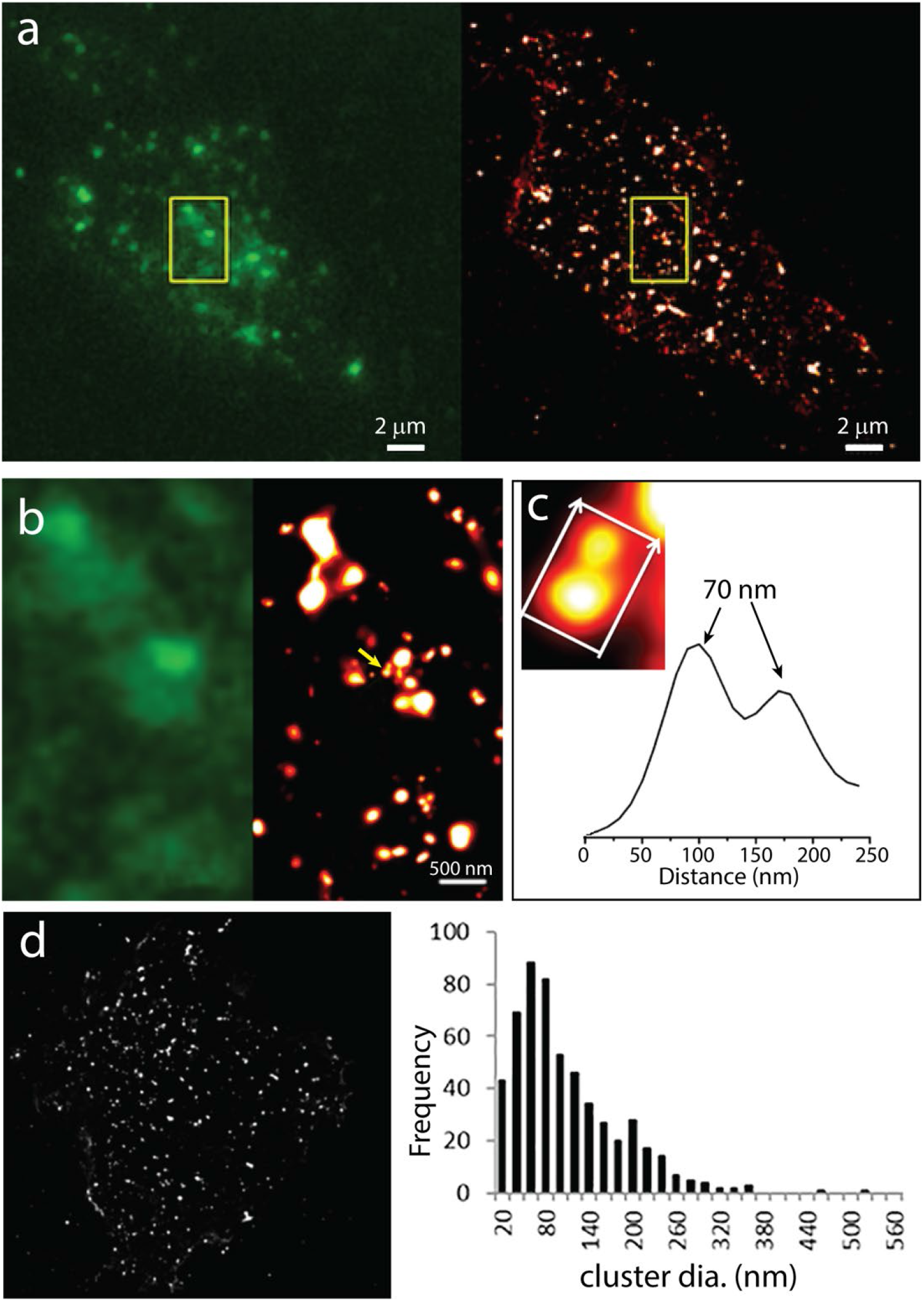
dSTORM images of AP3-Alexa 647 labeled TfR-EGFP cells. (a) Comparison of a diffraction limited TIRFM image of TfR-EGFP (left) and dSTORM reconstruction (right). (b) Zoomed image of the boxed region in (a). (c) Resolution analysis of the two TfR clusters pointed by the yellow arrow in (c) showing we can distinguish between two sub-resolution TfR clusters that were only 70nm apart. (d) The cluster size distribution of the corresponding reconstructed dSTORM image.

As show in the Figure 3, and similar to antibody labelling, the AP3-Alexa 647 aptamer is not cell permeable. It is however; significantly smaller than a conventional antibody and readily diffuses into cells after a mild detergent permeabilization step or other methods such as electroporation. To demonstrate labelling of intracellular proteins with AP3-Alexa647 aptamer for super-resolution imaging, we used a human fibroblast cell line (NIH 3T3) that express EGFP-lamin A, a structural protein located on the inner nuclear envelope (Fig. 4a). We developed a labeling protocol that uses a high-PEG buffer (“Buffer-S”) to stabilize nuclear structures while permeabilizing cells with a low concentration of Triton X-100. The permeabilized cells were incubated with AP3-Alexa 647 and the washed before PFA fixation. TIRFM and dSTORM imaging showed that AP3-Alexa 647 cleanly labelled the nuclear lamin A and the STORM reconstructions showed greatly improved resolution compared to the conventional TIRF image (Fig. 4b and c). We saw meshwork structures of lamin A (Fig. 4c) similar to what was previously reported in a super-resolution imaging study of nuclear lamin structures using structured illumination microscopy (SIM) (17), but at higher resolution (~30 nm) then the 2-fold improvement afforded using SIM. This also enabled us to visualize ring-like puncta structures on the nuclear membrane of many cells (Fig. 4c) that were uniform in size and about 1μm outer diameter with a central pore of about 300 nm. In another set of experiments, we co-stained the EGFP-lamin A cells with a red ER-tracker dye and imaged using confocal microscopy (Fig. 4d) and found numerous tubular structures that penetrated into and through the nucleus forming a structure known as the nucleoplasmic reticulum (18,19). There were large protrusions that were clearly visible by confocal imaging, but also numerous smaller structures which appeared as lamin “dots” in the confocal microscope (Fig. 4b,c left panel), but were found to have a clear ring-like structure in AP3-Alexa 647 dSTORM images (Fig. 4b,c right panel).

**Figure 4.**
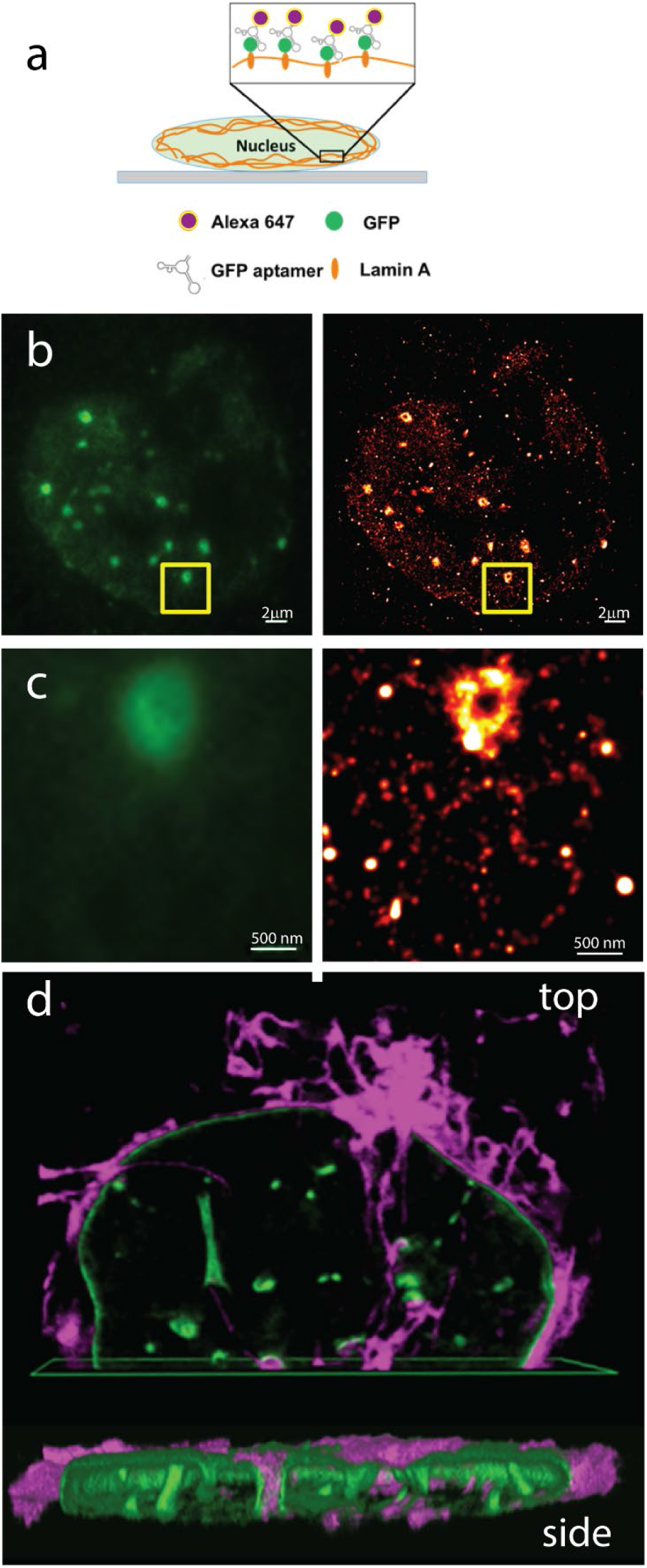
dSTORM images of AP3-Alexa 647 labeled Lamin A-EGFP cells. (a) Schematic of AP3-Alexa 647 labeled lamin A-EGFP cells. (b) dSTORM images of AP3-Alexa 647 labeled lamin A-EGFP cells and (c) zoomed image of the boxed region in (b). Cells were lightly permeabilized in an imaging buffer (“Buffer-S”) designed in maintain nuclear and organelle integrity and structure to allow for intracellular labelling. (d) 3D view of ER and lamin stained nucleus showing the extent and complexity of the nucleoplasmic reticulum. Using AP3-Alexa 647 dSTORM we found that in addition to the larger ER filled channels that protrude through the nucleus and are clearly visible in confocal microscope, there are numerous small protrusions where we visualize a ring/tunnel structure composed of lamin A. We take these to be small diameter protrusions on the ER into the nucleus that show up as small “dots” in the confocal image, but are sub-resolution ER tunnels in the nuclear lamina that are visible when imaged using dSTORM.

We interpret these to be smaller diameter “fingers” of the nucleoplasmic reticulum - ER protrusions into the nucleus are thought to be involved in signal transduction. These nuclear ER protrusions have been shown to contain functional IP3-mediated calcium signaling machinery (18) and are likely involved in the regulation of 3D chromatin structure and transcription. Interestingly, staining the cells with antibodies against the nuclear pore complex, we discovered that nuclear pore protein complexes also appeared in and around these tubular structures; however, over the surface of the nuclear lamina, there was no overall colocalization of nuclear pore protein complexes and the lamin meshwork (Supplementary Fig. S4).

### Super-resolution imaging with Aptamer-PAINT

Although dSTORM imaging provides the ability to resolve structures with lateral resolutions as high as 10-20 nm, the technique is primarily restricted to imaging fixed samples. Although there are reports of imaging live cells with STORM (16), the imaging conditions and buffer requirements are not well-suited for live-cell imaging. An oxygen scavenger system is required, the buffers have high concentrations of reducing agents, and high laser powers need to be used. More recently, a new localization based super-resolution technique called PAINT (Points Accumulation for Imaging in Nanoscale Topography) was developed (20, 21). Instead of using photo-switchable dyes, PAINT achieves molecular localization by having a pool of diffusing fluorescent molecules transiently bind to the target long enough so that on average, the location is brighter than the background and is therefore localizable. The most common PAINT method uses dye labeled ssDNA oligos (imager strands) with the sequences complimentary to the immobilized docking strand on the antibodies targeting proteins and/or structures of interest. Compared to dSTORM, PAINT has several advantages. It does not require the fluorophores to be photo-switchable, so there is a wider choice of fluorescent probes that can be used. Lower laser powers are used in PAINT and no harsh buffer conditions are required, and since the imager strands are replenished by diffusion, there is effectively no photobleaching. For these reasons, DNA-PAINT has potential as a live-cell super-resolution imaging technique.

A drawback of the conventional DNA-PAINT method is that is carried out using antibodies (20), and multiple staining steps are needed for attaching the docking strand. Here, we show that our aptamer-PAINT procedure is easier to implement since our modified AP3 aptamer already has the docking strand on one end making labeling a one-step process. To demonstrate aptamer-PAINT super resolution microscopy, we applied it in both fixed and live cell experiments. We designed a new version of AP3 with a 9nt linker on its 3’end that is complimentary to the sequence of an Alexa647 labeled ssDNA imager strand. The imager interacts with AP3 through DNA-RNA hybridization (Fig. 5a). We tested the binding of the AP3-docking sequence by conjugating it with Alexa 647 and used it to label TfR-EGFP expressing cells to ensure that the added linker on the 3’ end does not disrupt normal AP3 – EGFP binding. We then applied the aptamer PAINT method on fixed and permeabilized lamin A-EGFP expressing cells, and collected 50,000 images at a 50 Hz frame rate. Supplementary movie 1 shows the fluorescence blinking from transient interactions between AP3-docking strands and the diffusing imager strands. The reconstructed aptamer-PAINT images showed similar resolution improvement to what we achieved using dSTORM and AP3-Alexa 647 (Fig. 5b and 5c) revealing the same meshwork structures and ~micron diameter puncta on the nuclear envelope.

**Figure 5.**
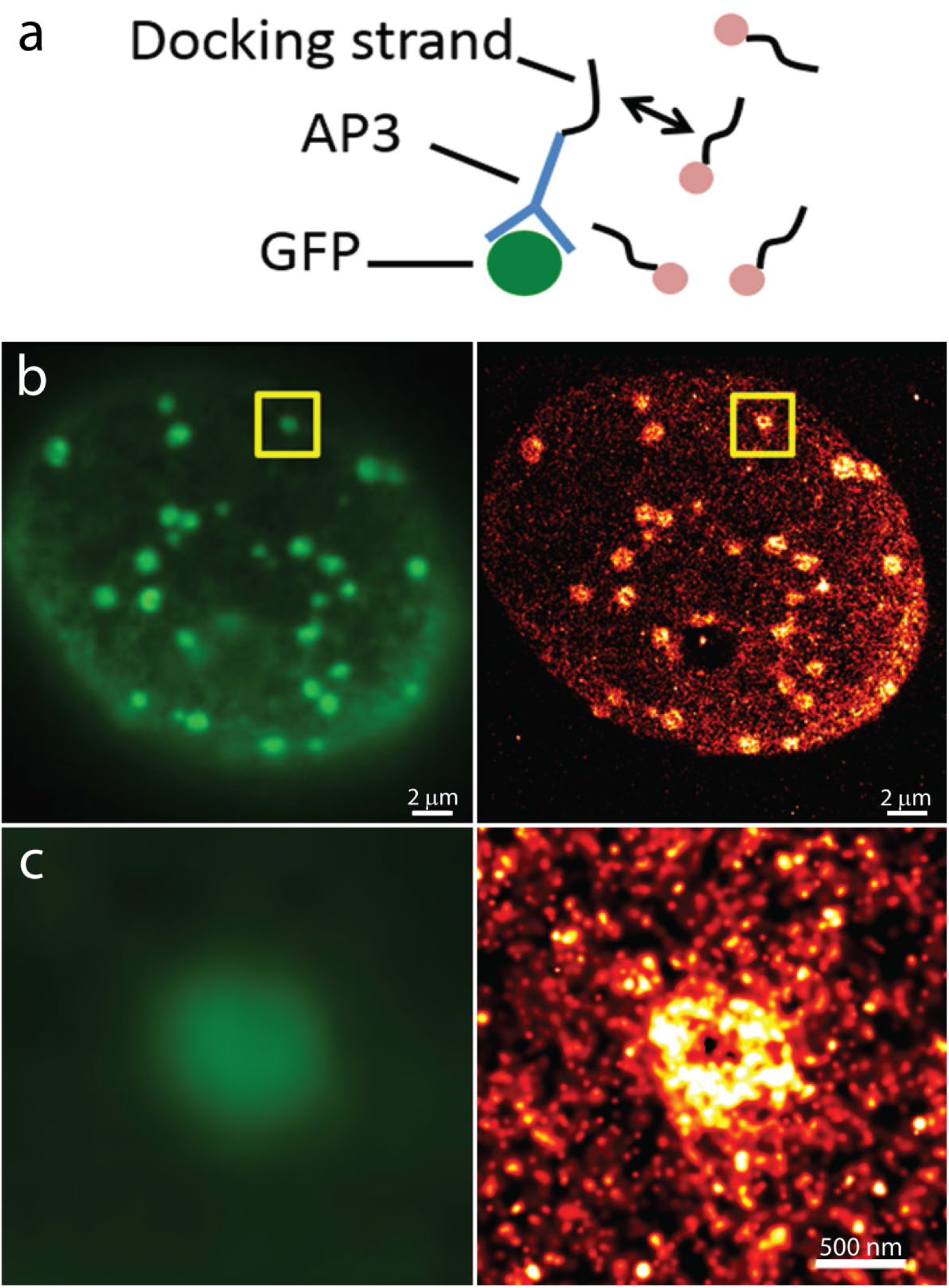
Aptamer PAINT images of AP3-Alexa 647 labeled Lamin A-EGFP cells. (a) Schematic of AP3-docking for PAINT super-resolution imaging. (b) TIRFM (top right) and PAINT reconstruction (bottom right) of lamin A-EGFP on the nuclear envelope, scale bar: 2um. (C). Zoomed-in boxed region of PAINT reconstruction (right) that shows the sub-diffraction limit lamin A structures, which is not shown on the corresponding TIRF image (left), scale bar: 500nm.

### Live-cell super-resolution imaging with Aptamer PAINT

Compared to dSTORM, where full power of the 642 nm laser in TIRF mode (0.5 mW measured in epi-mode) was required to drive fluorophore blinking, much lower intensity was used (2%-5%) for acquiring the aptamer-PAINT images. In addition to being live cell compatible, lower laser intensities also resulted in less background autofluorescence and therefore improved signal-to-background ratio (SBR), a parameter critical for localization based super-resolution methods. Unlike the buffer conditions used in dSTORM, the aptamer-PAINT buffer system is live cell imaging friendly. Taken together, these differences make aptamer-PAINT is an excellent candidate for super-resolution imaging applications in living cells where the high temporal resolution is not a requirement.

As a demonstration of live cell super-resolution imaging with aptamer-PAINT, we imaged the dynamics of TfR trafficking and recycling in the plasma membrane in EGFP-TfR expressing cells. HEK293 TfR-EGFP cells were plated on fibronectin coated MatTek dishes and images were acquired at 20 Hz. To examine TfR cluster dynamics, we created a movie comprised of reconstructed images from sequential 4000 frame segments of the aptamer-PAINT time series. Each composite image represents a “time-lapse” super resolution image of TfR clusters over a 3.33 min period. In contrast to SR images acquired from fixed cells, when imaging live cells we noticed that TfR on the outer membrane appeared as linear structures (Fig. 6a), which we interpret to be time-lapse traces of the receptor clusters being transported to endocytic sites. Figure 6b shows four frames from the first 13 minutes of the acquisition in which a cluster of ~75 TfR’s is seen moving towards a larger endocytic site. To verify that the cells were undergoing active receptor recycling and that the signal observed was solely from aptamer-PAINT labelled TfR-EGFP on the outer membrane, we removed the imager strands after incubating the AP3-docking strand-stained cells with them for 2-3 hours. We then added back Alexa 647 imager strands but saw no blinking, indicating the AP3-docking strand/TfR-EGFP complexes were now internalized and inaccessible to the imager strands. Actin filaments and motor proteins (Fig. 6d) are known to be involved with receptor recycling (22). To investigate their proximity and relationship to the linear arrays of TfR’s detected on the outer membrane, we stained actin in our TfR-EGFP cells with phalloidin-Alexa 647. Confocal images of TfR and actin are shown in Figure 6c. We found that the linear TfR structures on the outer membrane are colocalized with actin filaments inside the cell, possibly by linkage between the cytoplasmic tails of TfR and the motor proteins traveling along the actin filaments delivering the TfR clusters to the endocytic sites for recycling. Although our aptamer based PAINT is only applicable for capturing slower processes, it does provide the unique ability to quantify the number of proteins involved in a process such as receptor clustering and translocation to an endocytic site.

**Figure 6.**
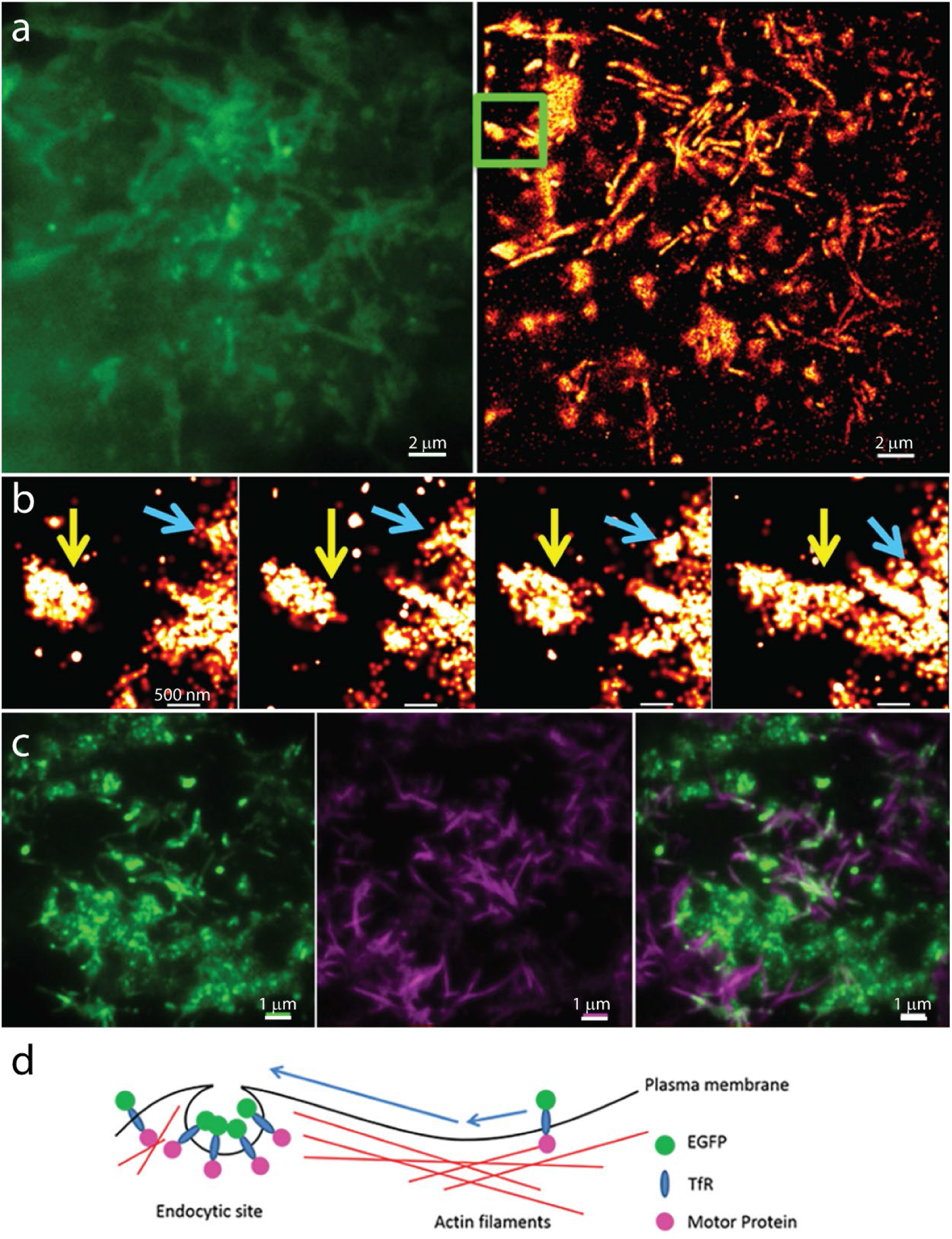
Aptamer PAINT images of AP3-Alexa 647 labeled live TfR-EGFP cells. (a) TIRF (left) and aptamer-PAINT reconstruction (right) of live HEK 293 TfR-EGFP cells. (b) Five consecutive PAINT reconstructions of the boxed region in (a) which show the dynamics of cluster trafficking on the cell membrane, as TfR’s are collect up and directed to endocytic sites (yellow and blue arrows). (c). HEK 293 TfR-EGFP cells stained with Phalloidin Alexa 647, which shows locations of actin filaments. They are spatially colocalized with tubular structures formed by TfR-EGFP. (D) A diagram showing that actin filaments act as a mediator for the transportation of TfR to the endocytic sites for recycling.

### Other advantages of RNA aptamer labeling

Nucleic acid aptamers have unique properties that can be exploited to extend their usefulness as fluorescent imaging labels. For example, aptamers can be easily removed by RNase treatment and reapplied with a different color fluorophore or at a different time. Sequential multiplexed imaging (23) uses this strategy to identify multiple different targets. Figure 7a shows two rounds of sequential binding of AP3-Alexa 647 to GFP beads. Simple modifications of the AP3 aptamer are easy to carry out and one can increase the per-probe brightness by adding an n-fold hybridization sequence that binds complementary labeled DNA’s. We added a 60nt linker at the 3’ end of AP3 that hybridizes with 3 labeled ssDNA oligos (Fig. 7b). Interestingly, a comparison of the average brightness of beads labelled with equal concentrations of AP3-Alexa 647 to our hybridized version that binds three dyes per aptamer, showed larger than expected signal increase (~4-fold). We speculate the increase above the expected 3-fold enhancement may be due to the absence of Alexa 647 quenching that may be occurring in the single fluorophore case, since the three fluorophores are structurally further away from the target site around which the AP3-Alexa 647 is tightly wrapped. Overall, the ability to easily design and employ hybridizable RNA probes of greater brightness or with different spectral codes (by mixing fluorophores), will be useful in a number of localization, tracking and spectral barcoding applications.

**Figure 7.**
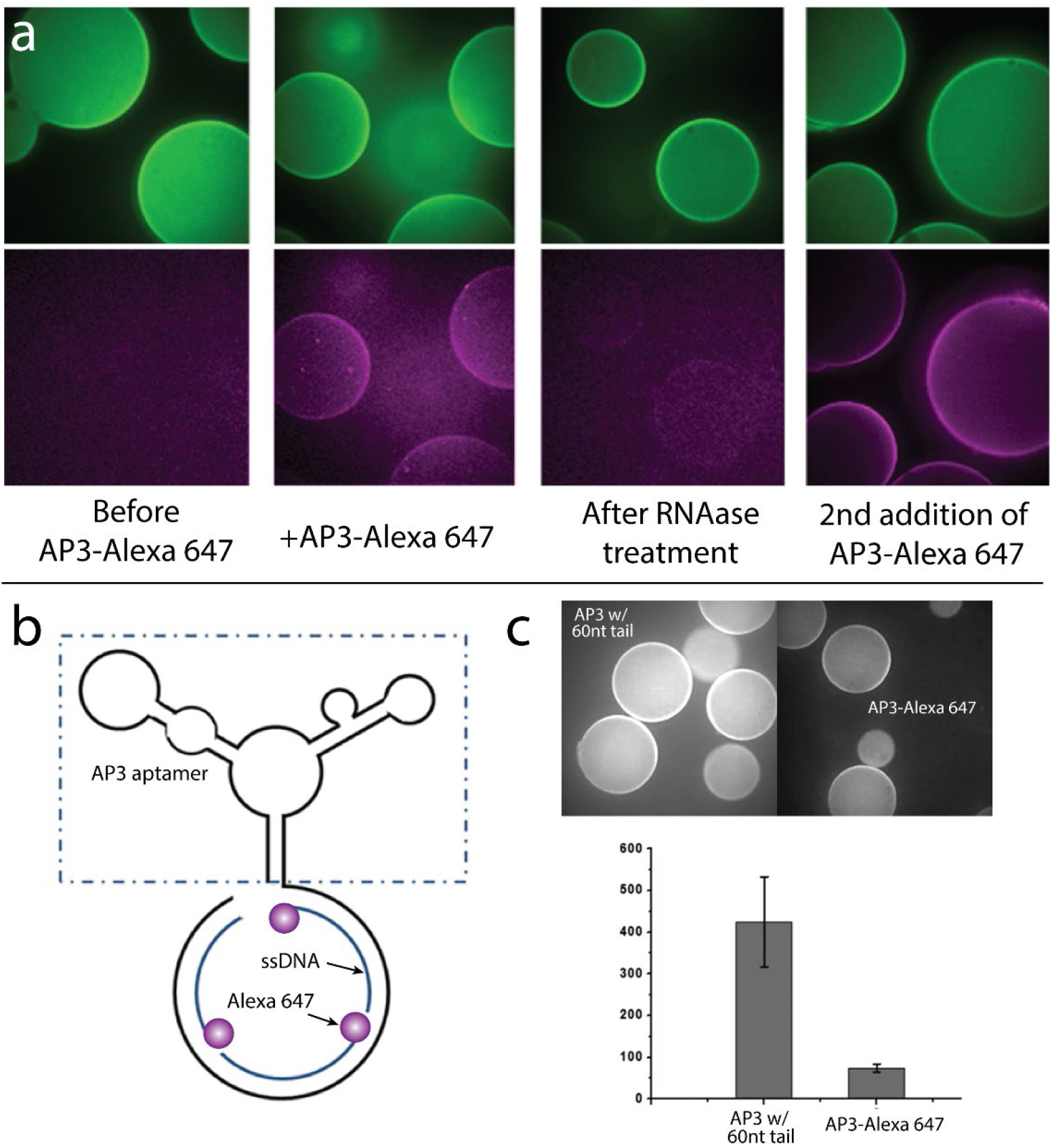
Additional unique advantages of aptamer labeling. (a) Two-rounds of sequential binding of Alexa 647 labeled AP3 on GFP beads using RNase treatment demonstrating aptamers as multiplexing reagents. (b) Left: structure of modified AP3 with 60nt linker at its 3’end that is hybridized to bind three labeled ssDNA oligos. Right: brightness comparison of beads with directly labeled AP3 and the 3X modified version (equal concentrations of AP3 were used).

## DISCUSSION

We have developed a novel super-resolution method based on using dye labeled RNA aptamers against a common protein tag, GFP, for imaging cellular structures on the plasma membrane as well as intracellular targets in permeabilized cells. The large number of GFP-tagged proteins available made the GFP/GFP-aptamer pair a logical choice for our proof-of-principle experiments. In some respects, aptamer-labelling is similar to antibody labelling, but aptamers are easily produced in the laboratory and physically smaller than conventional antibodies (an important attribute for localization-based SR microscopy). They are similar in size to nanobodies, but less expensive. Our method facilitates the use of dSTORM-compatible dyes for STORM imaging, or by the addition of a hybridization tail to the aptamer, DNA-PAINT microscopy. The aptamer–PAINT method is a simpler PAINT labeling technique since it avoids the multiple steps required in the original antibody-nucleotide version. As an alternative super-resolution technique, PAINT is more compatible with live cell imaging than dSTORM, and although slow (as is any localization-based SR method), we demonstrated use of the method to image receptor recycling at the single receptor level.

Tag-targeted RNA aptamers possess other unique advantages, such as the ability to multiplex fluorophores. Antibodies can have multiple fluorophores as well, but due to the non-specific labeling often used with antibodies, it is not a constant number per antibody. With aptamers the possibility exists to create integer brightness probes, or more useful, to easily create unique spectral bar codes by hybridizing different color fluorophores to the same aptamer. Finally, we showed that RNA aptamers could be removed from the target protein by a simple RNase treatment, which may have uses in multicolor imaging experiments or in live cell pulse-chase applications. A final use not demonstrated here, but feasible, is the use of fluorophore-aptamer labeling to identify cells of the interest in laser microdissection processing of tissue samples for mass spectroscopy analysis. This is often done with antibodies, but the presence of protein antibody adds complexity to the MS analysis, which is avoided by nucleic acid aptamer tags.

## DATA AVAILABILITY

The AP3 sequence is given in the Materials and Methods of this publication.

## Supplementary Figures

**Figure S1.**
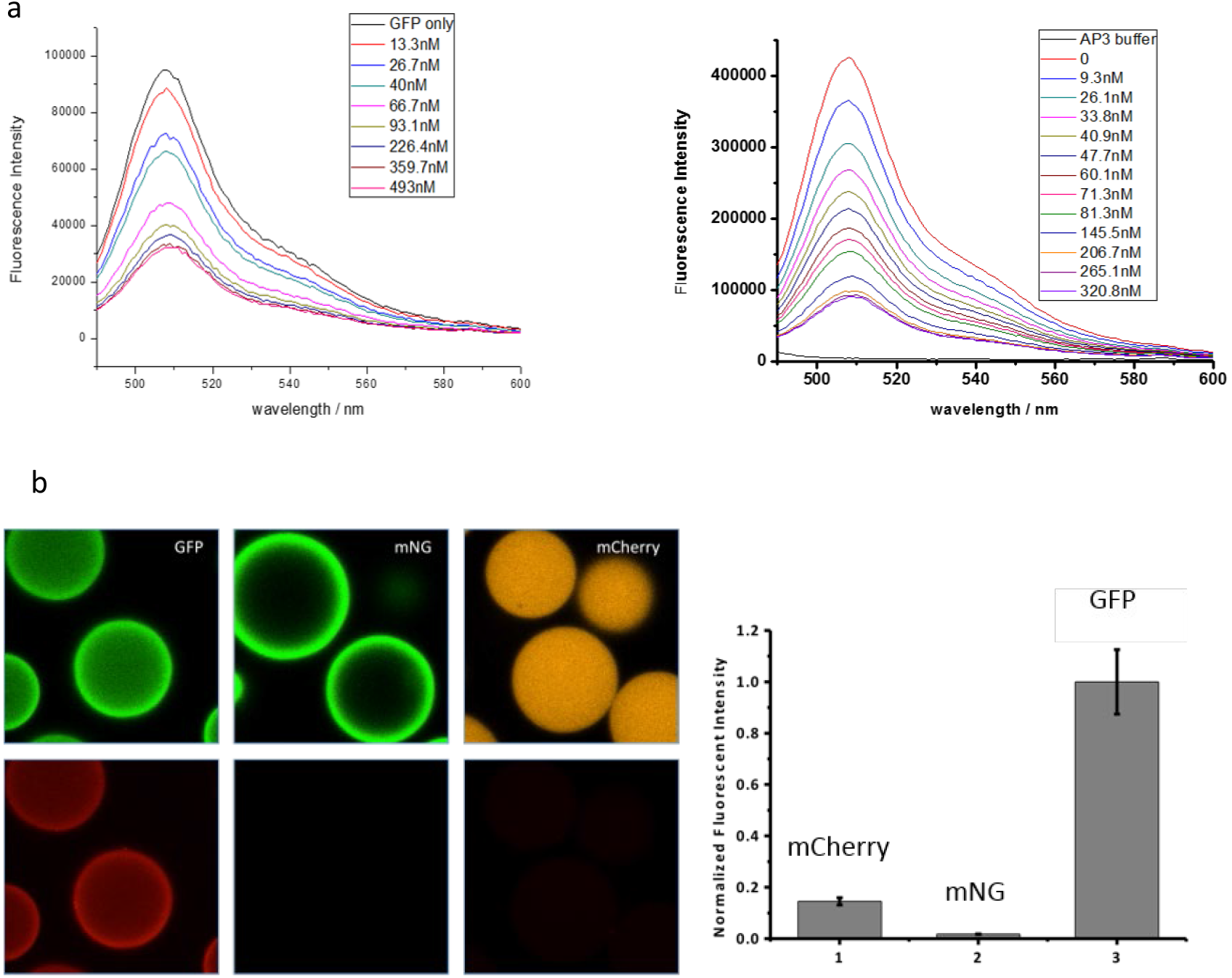
(a) The measured fluorescence intensity of EGFP decreases as increased number of EGFP molecules were bound by unlabeled AP3 (left) and Alexa 647 labeled AP3 (right). This data was used to calculate binding affinity Kd of both unlabeled and labeled AP3. (b) The binding tests of AP3-Alexa 647 to other fluorescent protein such as mNeonGreen and mCherry shows that AP3 is highly specific for EGFP and its derivatives.

**Figure S2.**
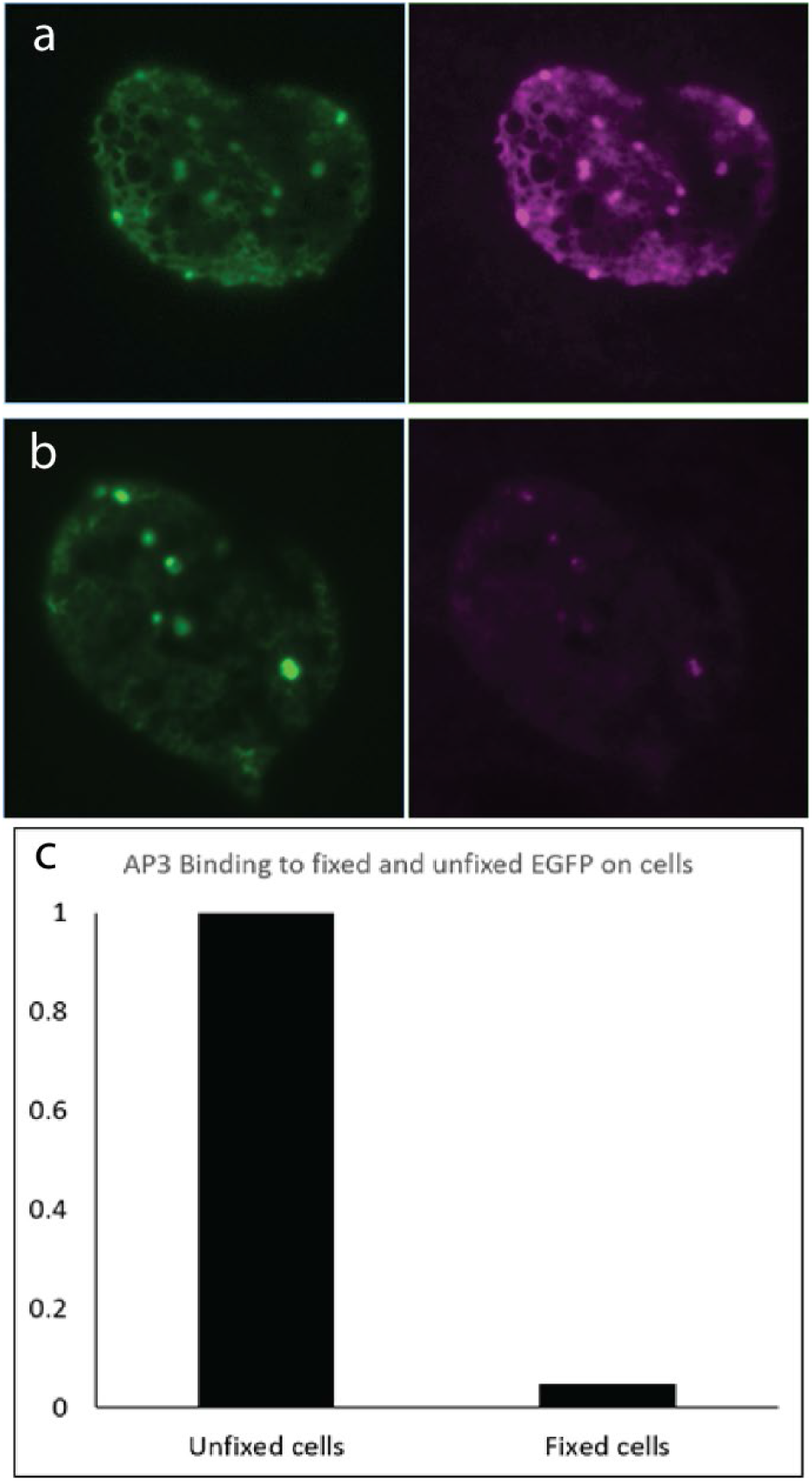
(a) AP3-Alexa 647 staining of live HEK 293 TfR-EGFP cells. (b) AP3-Alexa 647 staining of fixed HEK 293 TfR-EGFP cells. The staining was done on the same conditions for 1 hour at room temperature and imaged with the same laser power for both green and red channels. (c) Comparison of AP3-Alexa 647 binding to fixed and unfixed TfR-EGFP on cell’s plasma membrane. Background was subtracted and fluorescent signal of red channel was normalized to the green channel.

**Figure S2.**
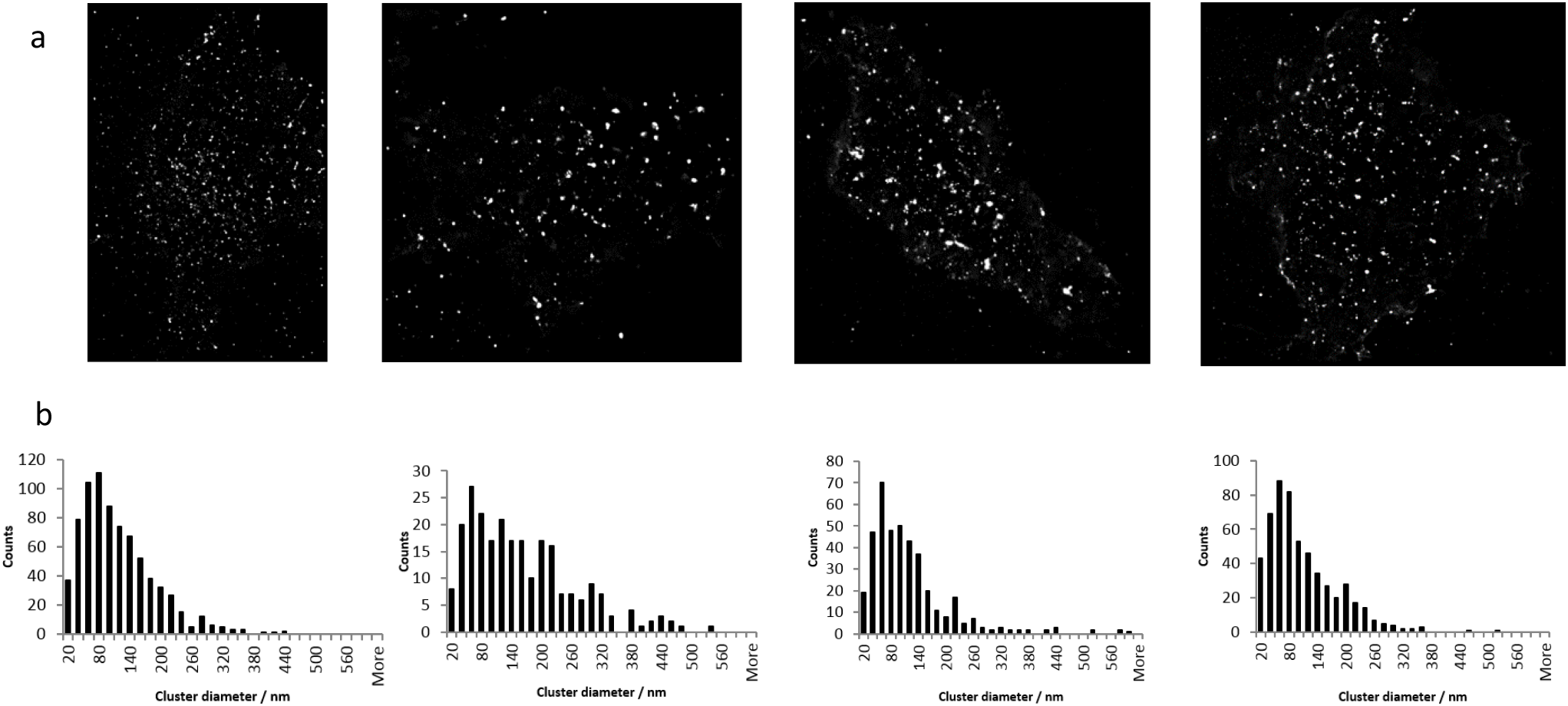
(a) Reconstructed dSTORM images of four distinct HEK293 EGFP TfR cells. (b) The cluster size distributions of the corresponding reconstructed dSTORM images in (a). The TfR cluster size distribution measured is in accordance with what has been reported in the literature, where the peak is between 60 and 80 nm in diameter.

**Figure S4.**
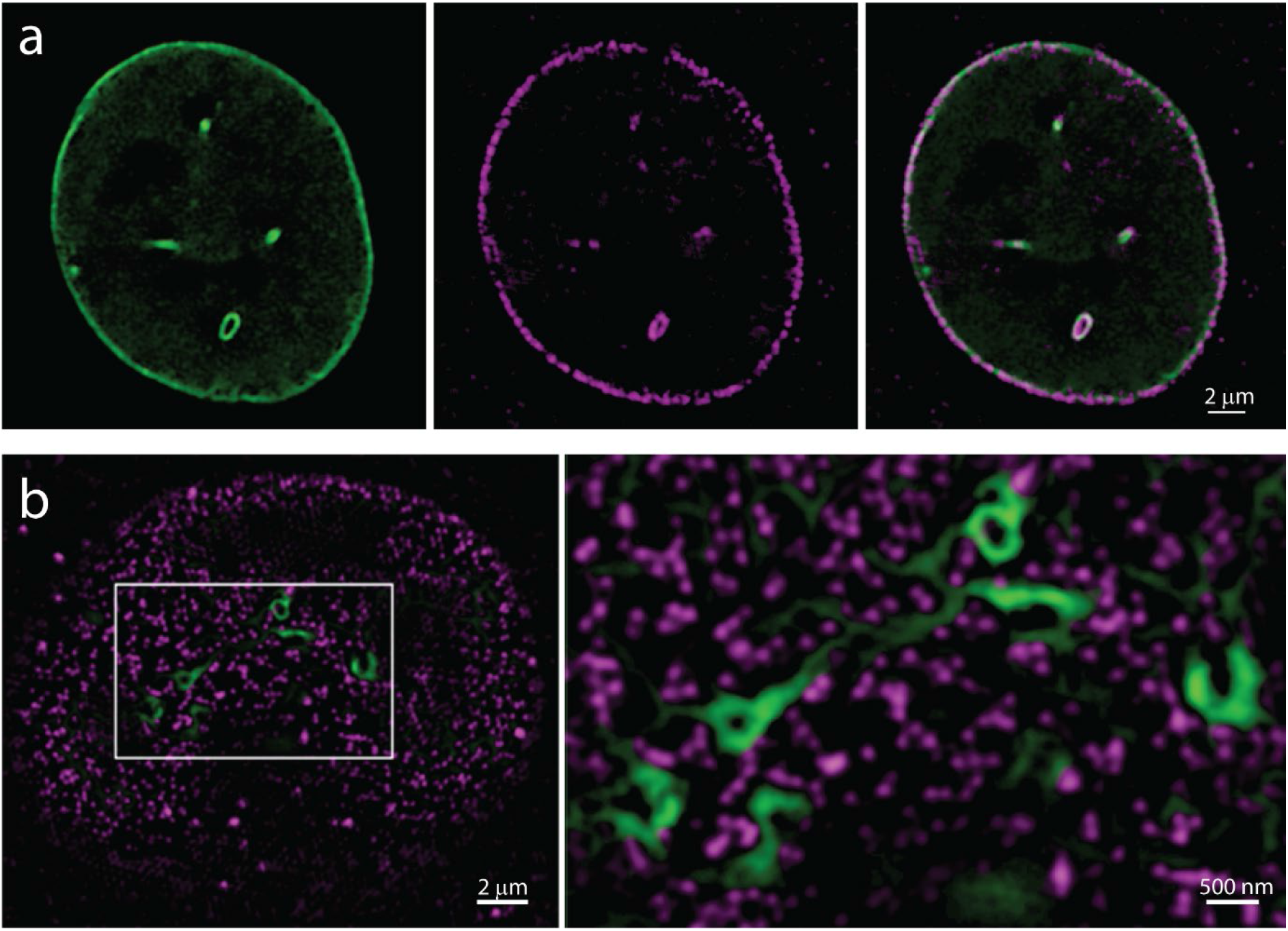
Super-resolution Structured Illumination Microcopy (SR-SIM) images of Nuclear Pore complex (NUP) antibody-Alexa 647 (red) stained Lamin A-EGFP (green) cells. (a) Optical section through the center of the nucleus showing that internally, nucleoplasmic reticulum protrusions have nuclear pore protein complexes associated with them. (b) On the nuclear lamina NUP complexes tend to be excluded from the Lamin A protein matrix.

